# Transcallosal generation of phase aligned beta bursts underlies TMS-induced interhemispheric inhibition

**DOI:** 10.1101/2024.11.22.624677

**Authors:** Christian Georgiev, Scott J. Mongold, Pierre Cabaraux, Gilles Naeije, Julie Duque, Mathieu Bourguignon

**Affiliations:** Laboratory of Functional Anatomy, Faculty of Human Motor Sciences, Université libre de Bruxelles (ULB), 1070 Brussels, Belgium; Department of Neurology, Hôpital universitaire de Bruxelles (HUB), Université libre de Bruxelles (ULB), 1070 Brussels, Belgium; Laboratoire de Neuroanatomie et Neuroimagerie translationnelles, UNI – ULB Neuroscience Institute, Université libre de Bruxelles (ULB), 1070 Brussels, Belgium; Centre de Référence Neuromusculaire, Department of Neurology, CUB Hôpital Erasme, Université libre de Bruxelles (ULB), 1070 Brussels, Belgium; Institute of Neuroscience, Université catholique de Louvain, 1200 Brussels, Belgium; WEL Research Institute, avenue Pasteur, 6, 1300 Wavre, Belgium

**Keywords:** beta oscillation, beta bursts, phase-alignment, interhemispheric inhibition, ipsilateral silent period, bimanual dexterity

## Abstract

The excitability of the sensorimotor (SM1) cortices is reflected in the bilateral ∼20 Hz beta oscillations. The extent to which these oscillations subtend the interhemispheric inhibition (IHI) captured by the Transcranial Magnetic Stimulation (TMS) ipsilateral Silent Period (iSP) protocol remains unclear. Therefore, we investigated the relationship between movement-related beta suppression and the iSP, along with their role for manual dexterity. Forty adults underwent an Electroencephalography assessment of beta suppression during volitional left hand movement and a TMS assessment of iSP recorded from the right hand. In both cases, left SM1 beta oscillations (contralateral to the activated right SM1), were monitored through a proxy signal – the Electromyography of the contracted right hand. Bimanual dexterity was assessed with the Purdue Pegboard. Volitional movement caused significant bilateral SM1 beta suppression in nearly all participants (≥ 85 %). ISPs were observed in every participant. In the proxy signal for the left SM1, the iSP coincided with TMS-evoked high-amplitude beta bursts. These bursts showed significant phase alignment across participants 10–70 ms after the TMS pulse. There was no significant association between the left-/right-hemisphere beta suppression, iSP, and bimanual dexterity. Our results highlight the distinct nature of beta oscillation changes during volitional movement compared to TMS-iSP and show that TMS induces IHI via transcallosal generation of phase aligned beta bursts. Furthermore, our data suggests that only the initial phase of a beta burst carries an inhibitory effect. It also highlights the possibility of evoking a beta burst with the iSP protocol, opening perspectives for future neuroimaging and modeling studies.

## 1. Introduction

Beta oscillations, comprised of non-rhythmic bursts of 13-30 Hz activity (Rayson et al., 2023; Shin et al., 2017; Szul et al., 2023), are commonly observed in Electroencephalography (EEG) and Magnetoencephalography recordings of the primary sensorimotor cortex (SM1; Pfurtscheller, 1981; Salenius and Hari, 2003; Neuper et al., 2006; Démas et al., 2020). Numerous studies have implicated the beta oscillations in the balancing of excitatory and inhibitory processes within the SM1 (Sherman et al., 2016; Khanna and Carmena, 2017). Particularly, a suppression of beta bursts reflects the neuronal activation within a given somatotopic region and an enhancement of beta bursts reflects the subsequent incoming neuronal inhibition (Heinrichs-Graham et al., 2017; Wessel, 2020). However, the ability of beta oscillations to synchronize across distant cortical areas (Buzsáki and Draguhn, 2004) has led researchers to investigate beta dynamics on a larger scale. This has provided consistent evidence that the beta suppression observed during bodily movement is typically bilateral (Pfurtscheller and Lopes da Silva, 1999; McFarland et al., 2000; van Wijk et al., 2012) and such bilaterality defines a sensorimotor beta-band network (Mary et al., 2017; Sugata et al., 2020). However, whether such large-scale bilateral beta oscillatory dynamics reflect the transcallosal regulation of excitation and inhibition between the two SM1s remains mostly unclear.

One mechanism theorized to realize transcallosal SM1 regulations is interhemispheric inhibition (IHI; Bologna et al., 2012; Ferbert et al., 1992; Duque et al., 2005, 2007; Giovannelli et al., 2009; Fleming and Newham, 2016). IHI is a process through which activation of one brain region selectively suppresses activation of its homologous region in the contralateral hemisphere (Perez and Cohen, 2009). IHI has been widely investigated with Transcranial Magnetic Stimulation (TMS) paradigms such as the ipsilateral Silent Period (iSP). The iSP manifests as a suppression of the Electromyography (EMG) activity of a contracted muscle after a TMS pulse is applied above the ipsilateral primary motor cortex (M1) representation of that muscle (Fling et al., 2011; Fleming and Newham, 2016; Giovannelli et al., 2009; Jung and Ziemann, 2006; Kuo et al., 2017). The TMS pulse inhibits the contralateral SM1 which disrupts its efferent output and results in the silenced contralateral (ipsilateral to the TMS pulse) EMG (Wassermann et al., 1991). Considering the strong implication of beta oscillations in cortical inhibition raises the question whether modulations of left-/right-hemisphere beta oscillations underpin the changes in M1 excitability captured by the TMS-evoked iSP.

Previous research with simultaneous TMS-EEG has shown that the potency of a TMS pulse is contingent on the amplitude, rate, and phase of naturally occurring beta bursts (Hussain et al., 2019). However, beta oscillations are highly variable across individuals (Illman et al., 2022), prompting the question whether this inter-individual variability results in inter-individual differences in IHI captured by the iSP. Furthermore, it is unclear exactly which feature of the beta bursts carries the inhibitory effect. Interestingly, single-pulse TMS above a certain brain region can alter the amplitude, as well as the phase of neuronal oscillations both within that given region (Premoli et al., 2017) as well as in another anatomically connected brain region (Kawasaki et al., 2014; Erickson et al., 2024). Therefore, it is possible that an iSP-evoking TMS pulse alters contralateral beta oscillations in a manner akin to the phase resetting theorized to underlie neural information transfer (Canavier, 2015; Voloh and Womelsdorf, 2016).

To elucidate the extent to which beta oscillations subtend IHI, we investigate the relationship between bilateral changes in beta amplitude during volitional movement and iSP magnitude. Subsequently, building on findings showing that SM1 beta oscillatory activity propagates to, and can be reliably assessed from peripheral muscle activity (Bourguignon et al., 2017; Bräcklein et al., 2022; Echeverria-Altuna et al., 2022; Mongold et al., 2022), we used the EMG of the contracted muscle during the iSP protocol as a proxy for assessing the effect of the TMS pulse on the contralateral cortical beta oscillations. Lastly, to explore the behavioral relevance of the beta oscillations and the iSP, we investigated their association with bimanual motor dexterity.

## 2. Materials and Methods

### 2.1 Participants

Forty (16 female; Mean ± SD age = 24.5 ± 4.4, range 20-49 years) adult volunteers with normal or corrected-to-normal vision and hearing participated in the study. All participants were right-handed according to the Edinburgh Handedness Inventory (Oldfield, 1971), did not suffer from neurological or psychiatric disorders, did not take any medication known to alter cerebrocortical excitability, and had no contraindications to TMS. The experimental protocol was approved by the ethics committee of the CUB Hôpital Erasme (CCB, B4062023000234) and the experiment was conducted in accordance with the Declaration of Helsinki. All participants signed an informed consent prior to participation.

### 2.2 Experimental Design

The experimental protocol was divided into three parts: an EEG assessment of cortical beta oscillations during volitional hand movement, a TMS assessment of IHI via the iSP protocol, and a behavioral assessment of bimanual dexterity. The three parts of the experiment were conducted in a randomized order within a single session. All participants successfully completed the experiment and no adverse effects of TMS were reported.

#### EEG assessment of the cortical beta oscillations

Participants’ task was to maintain a stable isometric pinch-grip contraction with the right thumb and index finger against a force transducer (RS Pro, P/N 1004-0.6 kg, Vishay Precision Group, Malvern, PA, USA) at 10 ± 3 % of their maximum voluntary contraction (MVC) force. MVC was individually determined for each participant prior to the start of the experiment. Task compliance was ensured with real-time visual feedback prompting participants to increase their force when it subceeded 7 % MVC and to decrease their force when it exceeded 13 % MVC. While engaged in the isometric contraction with the right hand, once every 3–4 s (random jitter), participants were presented with an auditory tone prompting them to perform a fast closing/opening movement with the fingers of their non-contracted left hand (Figure 1A). This task was performed for 2 blocks of 5 minutes with a short break in between.

**Figure 1.**
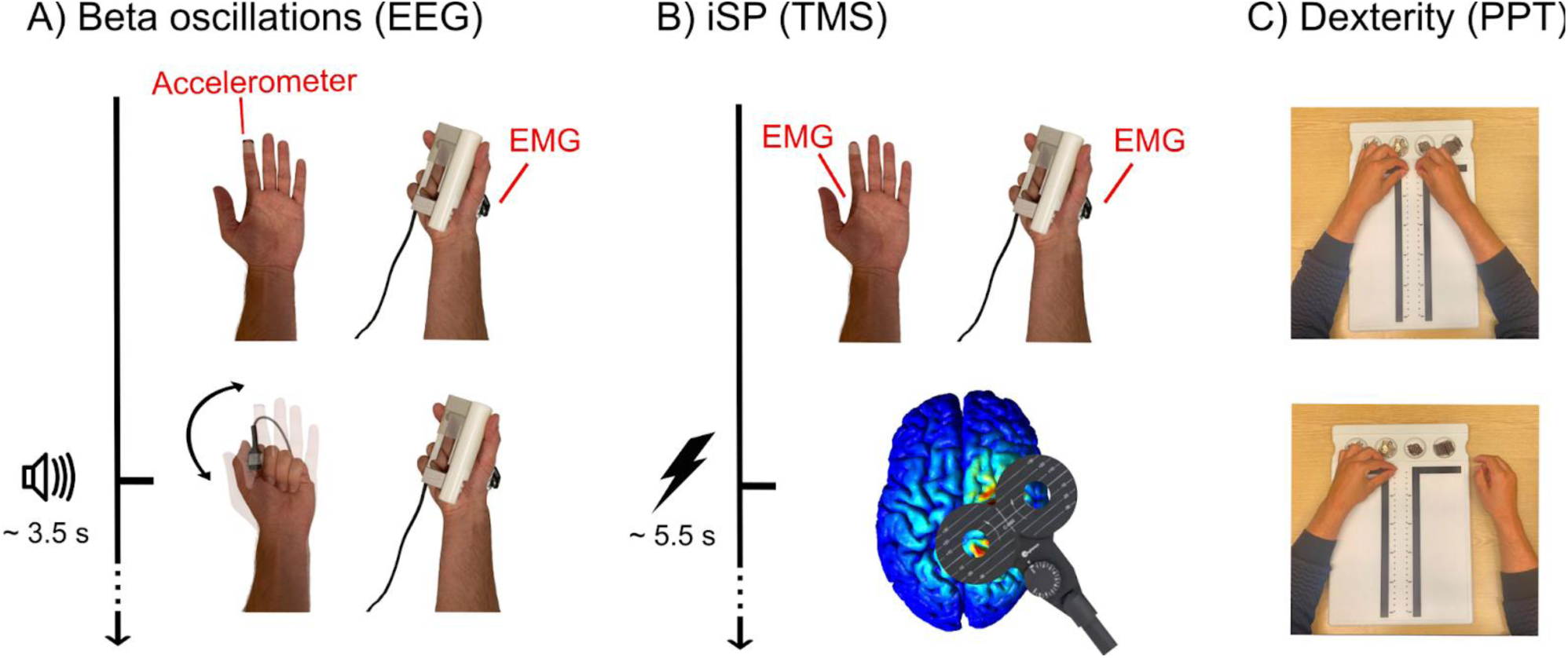
Experimental protocol. For assessing Electroencephalography (EEG) beta oscillations, participants held a constant contraction with their right hand monitored with Electromyography (EMG), and performed left hand movements in response to auditory prompts (A). For assessing interhemispheric inhibition, participants underwent an ipsilateral SIlent Period protocol with a constant contraction with their right hand monitored with EMG and single-pulse Transcranial Magnetic Stimulation (TMS) applied above the ipsilateral primary motor cortex (B). For assessing bimanual dexterity, participants performed the Purdue Pegboard Test (PPT) (C).

Throughout the two blocks, EEG was recorded with 64 Ag/AgCl-sintered electrodes arranged according to an augmented 10-20 system (EEGO Mylab, ANT Neuro, Hengelo, the Netherlands). Electrode impedance was kept blow 20 kΩ. The recording was online referenced to electrode CPz, and sampled at 1 kHz. The signals of a 3-axis accelerometer (ADXL335, AnalogDevices, Wilmington, MA, USA) attached to the index finger of the closing/opening left hand were recorded in synchrony with the EEG signals.

Surface EMG (COMETA Pico, COMETA, Milan, Italy) from the contracted right First Dorsal Interosseous (rFDI) muscle was recorded with a sampling frequency of 1 kHz. EMG was recorded in a monopolar configuration, with an “active” electrode placed above the rFDI muscle belly, and a “reference” electrode placed above the first metacarpal bone. The EEG and EMG acquisitions were synchronized via the delivery of digital synchronization triggers.

#### TMS assessment of the iSP

Participants were seated on a TMS chair and equipped with a neck brace for securing the head into position and minimizing unwanted movements. Surface EMG from the left and right FDI muscles were recorded as in the EEG task.

The hotspot corresponding to the hand area on the right M1 was identified via biphasic single-pulse TMS (MagPro X100, MagVenture, Farum, Denmark) with a hand-held figure-of-eight coil (7.5 cm external loop diameter), pointed 45° away from the midline. Each participant’s Resting Motor Threshold (RMT) was identified. RMT was defined as the minimal TMS intensity that could elicit a motor-evoked potential of at least 50 μV on 5 out of 10 stimulations (Rossini et al., 1994).

During the iSP task, participants held an isometric contraction with their right hand while completely relaxing their left hand. The parameters of the contraction and the visual feedback were identical to the ones during the EEG task, therefore allowing for direct comparison of right FDI EMGs between the two tasks. While participants were holding the contraction, once every 5–6 s (random jitter) a TMS pulse was delivered to the previously identified hotspot with an intensity of 130 % RMT (Figure 1B). The procedure was performed for 2 blocks of 4.5 minutes, resulting in a total of 100 iSPs per participant.

#### Behavioral assessment of unimanual and bimanual dexterity

Participants completed the Purdue Pegboard Test (PPT; Tiffin and Asher, 1948). The PPT assesses the fine motor performance of the left hand (*PPT_LH_*), right hand (*PPT_RH_*), both hands in synchrony (*PPT_BH_*), and both hands in alternation (*PPT_Assembly_; Figure 1C*). Before each subtest, the task was explained to the participants and they were allowed one practice trial. All scores were computed following the PPT guidelines (Tiffin and Asher, 1948). Two additional bimanual coordination scores were computed. The first coordination score (*PPT_C1_*) was the *PPT_Assembly_* divided by the geometric mean of the *PPT_LH_* and *PPT_RH_*. The second PPT coordination index (*PPT_C2_*) was the *PPT_Assembly_* divided by the *PPT_BH_*.

### 2.3 EEG and EMG Data Processing

#### EEG preprocessing

EEG data was preprocessed offline with a custom MATLAB (Mathworks, Natick, MA) script and the FieldTrip toolbox (Oostenveld et al., 2011). Bad channels were identified following the guidelines of Bigdely-Shamlo et al. (2015) and topographically interpolated. EEG signals were re-referenced to a common average, filtered through 0.3–45 Hz, and Independent Components Analysis (Vigário et al., 2000) was used for further artifact suppression. Twenty independent components were evaluated from the data with a FastICA algorithm with dimension reduction, 25; nonlinearity, tanh (Hyvärinen and Oja, 2000; Vigário et al., 2000); and independent components corresponding to eye-blink, eye movement, and heartbeat artifacts were visually identified and removed. On average, 4 (SD = 1.3) independent components per participant were removed.

#### Beta amplitude modulation analysis

EEG signals were filtered through 5-Hz-wide frequency bands centered on 5–40 Hz by steps of 1 Hz. The bandpass filter used to this aim was designed in the frequency domain with zero-phase and 1-Hz-wide squared-sine transitions from 0 to 1 and 1 to 0 (e.g., the filter centered on 20 Hz rose from 0 at 17 Hz to 1 at 18 Hz and ebbed from 1 at 22 Hz to 0 at 23 Hz). This filtering approach is standardly applied to electrophysiological data and allows for resolute time-frequency decompositions of the signals (e.g., Bourguignon et al., 2017; Mongold et al., 2022). Then, EEG signals were smoothly set to 0 (squared-cosine transition of 1 s) at timings 1 s around artifacts (time points where full-band EEG amplitude in at least one electrode exceeded 5 SDs above the mean), and to avoid edge effects, further analyses were based on time points at least 2 s away from artifacts and appearance of force visual feedback. Subband signals’ envelope was then extracted with the Hilbert transform and downsampled to 20 Hz. To avoid contamination by very slow envelope variations, these subband envelopes were divided by their low-pass filtered version at 0.1 Hz (squared-sine transition from 0.05 to 0.15 Hz). Such normalization ensures that envelopes fluctuate about the value 1, and are blind to changes slower than 0.1 Hz that cannot be ascribed to the task. As a further normalization, the envelopes were centered on 0 by subtracting the value 1, and converted to percentage of change by multiplying by 100 %.

Subband envelopes were segmented into epochs from –1000 to 2500 ms relative to left hand movement onset as identified in accelerometer recordings. Epochs which contained time points less than 2 s away from artifacts or in which participants failed to perform a movement or failed to maintain the force of the right hand at 10 ± 3 % MVC were excluded from the analysis. The remaining epochs (113 ± 25, Mean ± SD across participants) were subsequently averaged across trials, giving rise to a time-frequency map of amplitude modulation for each EEG channel and each participant. Further analyses were focused on preselected electrodes spanning the left (C1, C3, C5, CP1, CP3, CP5, FC1, FC3, FC5) and right (C2, C4, C6, CP2, CP4, CP6, FC2, FC4, FC6) SM1s. We further identified within each SM1 electrode preselection the electrode with the maximal amplitude suppression in a preselected time-frequency window ranging from – 500 to 1000 ms relative to left hand movement onset and from 13 to 30 Hz (βdepth). Within the corresponding electrode, we further extracted the size of the cluster of significant beta suppression in the previously defined time-frequency window (βsize; see subsection Statistical Analyses). Lastly, to highlight left-/right-hemisphere imbalance in beta modulations, contrasts between the left and right indices of beta suppression were computed as their difference (left minus right) divided by their sum, leading to Δ*β_size_* and Δ*β_depth_*. These contrasts are smaller than 0 when beta amplitude is more suppressed in the right hemisphere (contralateral to the moving hand) compared with the left hemisphere (ipsilateral to the moving hand) and vice versa.

#### EMG iSP preprocessing and analysis

The EMG data from the contracted rFDI during the iSP protocol was preprocessed offline with a custom MATLAB script. Data were filtered between 20 and 295 Hz, followed by notch filters at 50 Hz and its harmonics, and rectified. The EMG was segmented into epochs from –100 to 200 ms relative to TMS pulse onset. The epochs were averaged within participants and normalized by their baseline mean (taken from 50 ms to 5 ms prior to the TMS pulse), and used for iSP quantification. The iSP onset and offset were identified with Mean Consecutive Difference thresholding (Garvey et al., 2001; Hupfeld et al., 2020) and used for computing iSP latency (the time between the TMS pulse and the iSP onset), iSP duration (the time between the iSP onset and offset), and normalized iSP area (the integral of the normalized rectified EMG signal between the iSP onset and offset (Hupfeld et al., 2020)).

#### Time-frequency domain EMG analyses

To further characterize the beta modulations during the voluntary movements and the TMS-iSP protocol, we used the unrectified rFDI EMG of the contracted right hand during each task as a proxy for the changes occurring in the contralateral SM1 (Bourguignon et al., 2017; Bräcklein et al., 2022; Echeverria-Altuna et al., 2022; Mongold et al., 2022). EMG signals were filtered in subbands for envelope extraction, normalized by their <0.1 Hz trend, segmented, and averaged as previously done with the EEG signals. In addition, to separate the contribution of evoked and induced responses, the same analysis was also conducted on EMG signals from which the average response was subtracted.

#### EMG beta burst analyses

We further characterized the bursts underlying the beta modulations identified in the unrectified rFDI EMG during the volitional left hand movements and the TMS stimulations (Szul et al., 2023; Rayson et al., 2023). To this end, individual beta bursts in the EMG data from each task were identified following the methodology of Tinkhauser et al. (2017) and O’Keeffe et al. (2020). Beta envelopes were obtained using the Hilbert transformation applied to the unrectified EMG signals filtered through a 17-Hz-wide frequency band centered on 13–30 Hz. This filtering approach is commonly applied to electrophysiological data in preparation for single burst extraction (e.g., O’Keeffe et al., 2020). The EMG envelopes were further normalized by their <0.1 Hz trend. We then identified, for each participant and condition separately, time intervals where the EMG envelope exceeded its 75^th^ percentile. Such intervals were considered as beta bursts when their duration was greater than 50 ms. The time of occurrence of each beta burst was defined as the middle of the corresponding time interval. The probability of beta burst occurrence and the average burst duration were computed within 200-ms windows spaced by 100 ms and centered on -900 to 2400 ms relative to movement or TMS onset.

#### Phase domain EMG analysis

Lastly, having identified TMS-evoked beta bursts in the proxy signal for the left SM1, we further investigated whether TMS-evoked iSP relates to a phase resetting of these beta bursts in the rFDI EMG. Thus, the unrectified EMG was filtered as done for the beta burst analyses. Then, the EMG data was segmented into epochs from –200 to 500 ms relative to TMS pulse onset. Epochs were normalized by their root-mean-square amplitude (taken from 50 ms to 5 ms prior to the TMS pulse) and averaged across trials. Hilbert transformation was applied to the averaged EMG trace of each participant. Then, we determined whether Hilbert coefficients at time points from 0 to 200 ms by step of 10 ms tended to show a consistent phase across participants. Due to the uncertainty regarding EMG polarity (which is dependent on the choice of the active and reference electrodes that was not standardized across participants), phase consistency was assessed using an approach based on singular value decomposition (SVD). More precisely, for each time-point, SVD was applied to all participants’ Hilbert coefficients arranged as a matrix with the real part of all participants’ coefficients stacked as the first column and the imaginary part as the second column. In this setup, the SVD identified potential Hilbert coefficient alignment in the complex plane, corresponding to a phase alignment across participants of the EMG beta oscillations after the TMS pulse. The first singular value of the decomposition was used to quantify the strength of the phase alignment across participants.

### 2.4 Statistical analyses

#### Movement-related EEG and EMG beta suppression

Non-parametric cluster-based tests were applied to each participant’s EEG data to assess the significance of *β_size_* in the time-frequency window ranging from –500 to 1000 ms relative to left hand movement onset and from 13 to 30 Hz separately separately for the left (ipsilateral) and right (contralateral) hemisphere electrodes. In brief, 10,000 surrogate amplitude modulation maps were built based on EEG subband envelopes segmented with respect to random onsets separated by 3–4 s. These surrogate maps served to identify *β_size_*, for the genuine and each surrogate data, which was the size of the largest cluster of time-frequency bins (bin size: 50 ms × 1 Hz) displaying a larger amplitude suppression or enhancement than their percentile 95 across all maps. The *p*-value for *β_size_* was obtained as the proportion of surrogate *β_size_* values that were higher than the observed genuine value.

Analogous non-parametric tests were applied to the time-frequency maps of EMG amplitude.

#### Movement-related and TMS-evoked changes in beta burst properties

The statistical significance of changes in beta burst properties (probability of occurrence and duration) in each time interval from 0 ms to 500 ms relative to movement or TMS onset was assessed with a *t*-test in comparison with baseline values (mean from –900 to –500 ms). Correction for multiple comparisons was performed using Bonferroni.

#### TMS-evoked EMG beta phase reset

Surrogate-data-based statistics were applied to evaluate the significance of the TMS-evoked phase alignment in the beta band of the right FDI EMG. To do so, the strength of the phase alignment across participants for each time point after the TMS pulse was compared to a null distribution obtained using the Hilbert coefficients with a phase randomly selected in the range [-π, π] (10,000 repetitions) while preserving the amplitude. The *p*-value was obtained as the proportion of values in the null distribution that were higher than the genuine singular value.

#### Association Analyses

Redundancy analysis (RDA) was used to investigate the associations between all constructs of interest. RDA is a multivariate technique for modeling the effect of a linear combination of a set of related predictors on the linear combinations of a set of related outcomes (van den Wollenberg, 1977). In two RDA models, we assessed i) the predictive power of all indices of left-/right-hemisphere beta suppression imbalance on all indices of iSP and ii) the predictive power of all indices of left-/right-hemisphere beta imbalance and iSP together on all indices of bimanual motor dexterity. Statistical significance of each model was assessed with RDA Monte Carlo permutation tests with 10,000 permutations.

The RDA was followed-up with pairwise Spearman correlations between all variables of interest.

## 3. Results

### 3.1 Movement-related beta suppression

The EEG analyses identified a large-scale suppression of the beta amplitude following left hand movement (Figure 2). More quantitatively, in the time interval from –500 to 1000 ms, *β_size_*, was significant in 35 (87.5 %) participants (all *p* < 0.05) in the contralateral (right) hemisphere, and in 34 (85 %) participants (all *p* < 0.05) in the ipsilateral (left) hemisphere. Across participants, the contralateral *β_size_* was 100.5 ± 85.9 (Mean ± SD) and the ipsilateral *β_size_* was 84.4 ± 73.3 time-frequency bins, with the contralateral *β_size_* being significantly greater than the ipsilateral *β_size_* (*t*_39_ = 2.4, *p* = 0.024, *d* = 0.4). Likewise, across participants, the contralateral *β_depth_* was 26.1 ± 6.2 % suppression and the ipsilateral *β_depth_* was 24.1 ± 4.8 % suppression. The contralateral *β_depth_* was also significantly greater than the ipsilateral *β_depth_* (*t*_39_ = 4.4, *p* < 0.0001, *d* = 0.7).

**Figure 2.**
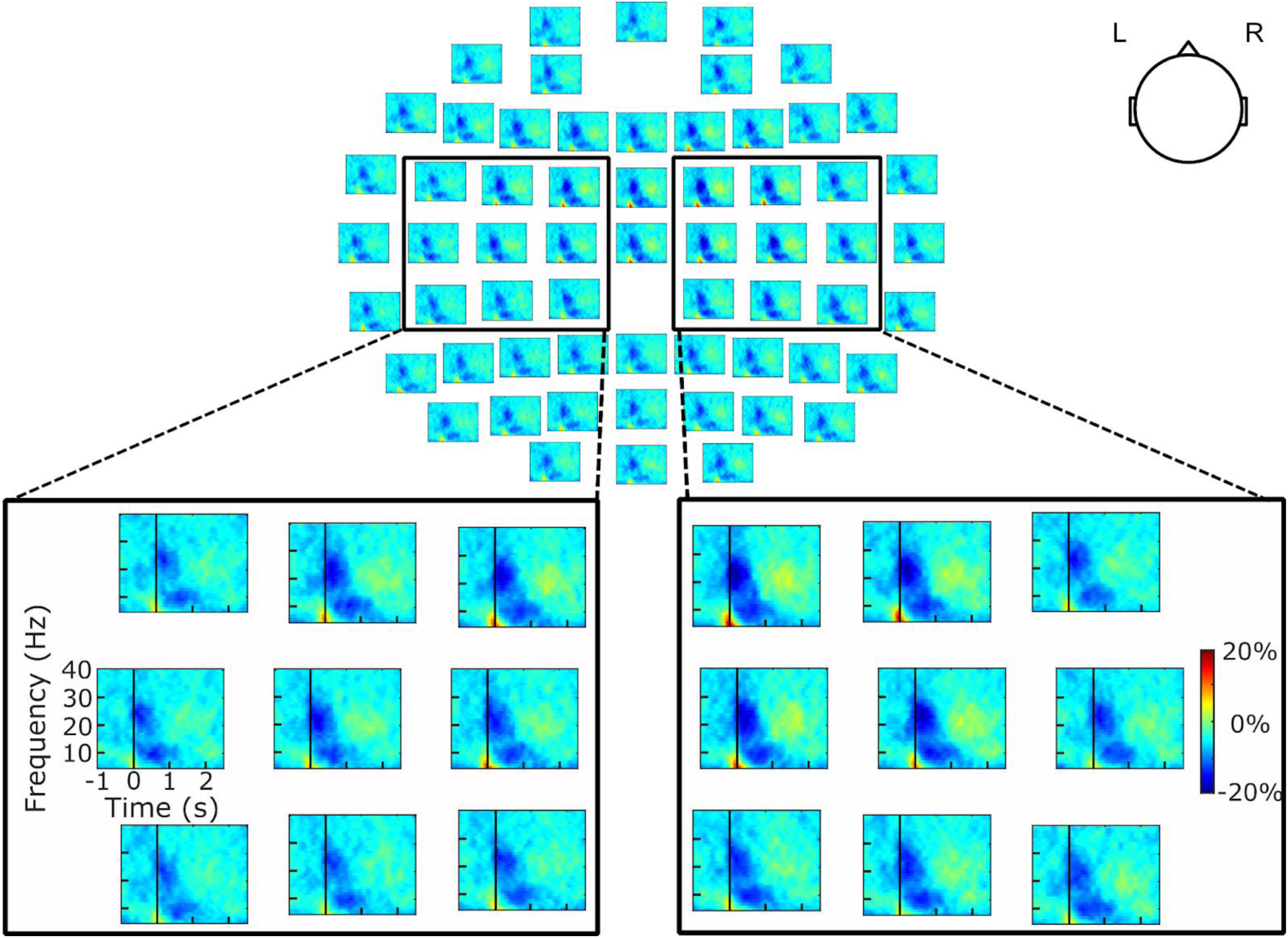
Group-averaged Electroencephalography scalp distribution of the time-frequency maps of amplitude modulation during the movement execution task. The black rectangles outline the data for the 9 electrodes of interest above each hemisphere’s primary sensorimotor cortex and the black vertical lines at time 0 indicate the onset of left hand movement as identified in accelerometer recordings.

### 3.2 TMS-evoked iSP

The RMT of our sample was 50.0 ± 8.6 % of the maximal stimulation intensity of the TMS device.

An iSP was identified in all participants (Figure 3), in the form of transiently decreased muscle activity ∼50 ms after TMS stimulation, characterized by a latency of 39.3 ± 6.2 ms, a duration of 25.1 ± 7.6 ms, and a normalized area of 7.1 ± 3.5 mV ms. Of note, iSPs were closely followed by two successive peaks of increased muscle activity up until ∼150 ms.

**Figure 3.**
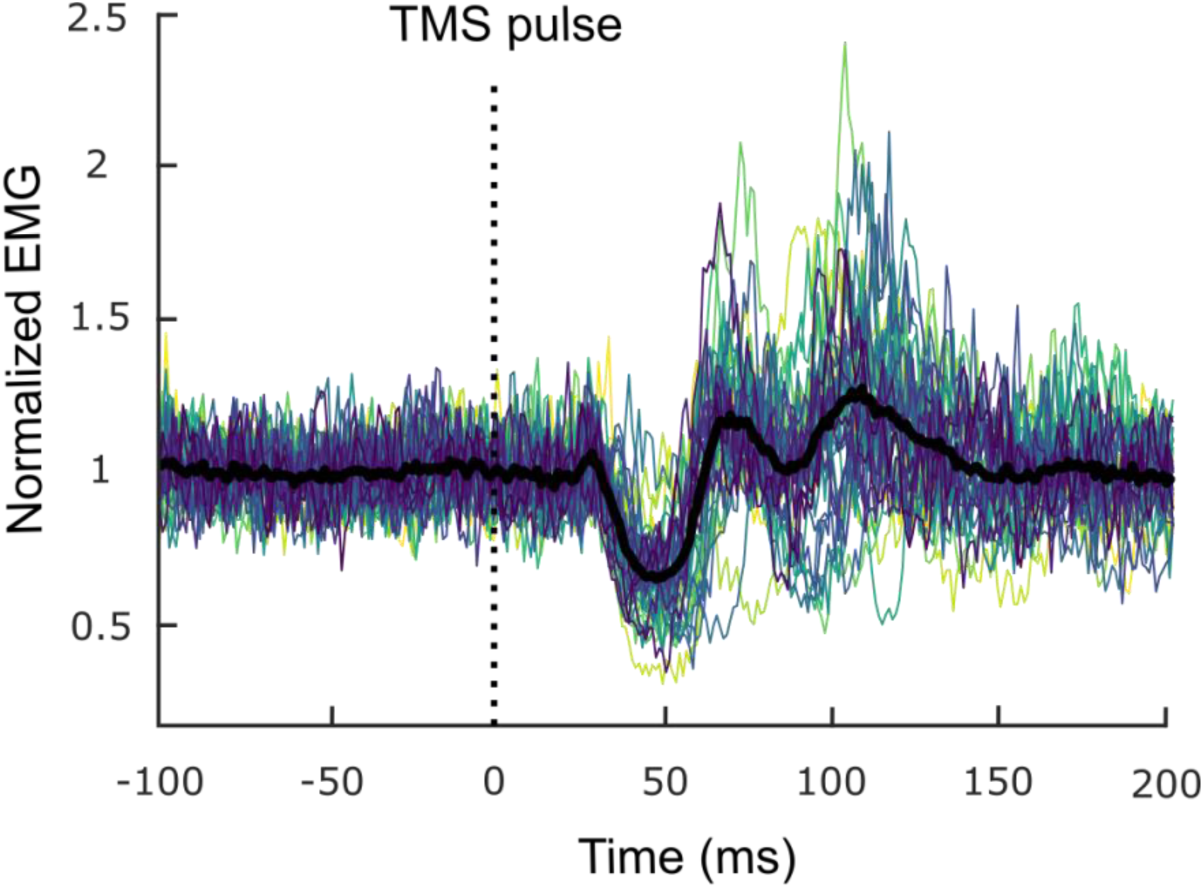
Normalized rectified Electromyography (EMG) traces (averaged across trials) during the ipsilateral Silent Period task. There is one trace per participant and the black trace corresponds to their grand average. The vertical dotted black line indicates the Transcranial Magnetic Stimulation (TMS) pulse onset.

### 3.3 Time-frequency domain EMG changes

Figure 4A presents the group-averaged time-frequency map of the amplitude of the rFDI EMG signal in the left hand movement task. It clearly highlights a large-scale suppression of the beta amplitude corresponding to the onset of the left hand movement, reflecting that observed in the left SM1 (Figure 2). In the time interval from –500 to 1000 ms, EMG *β_size_* was 53.2 ± 60.9 time-frequency bins and was statistically significant in 29 (72.5 %) participants (all *p* < 0.05).

**Figure 4.**
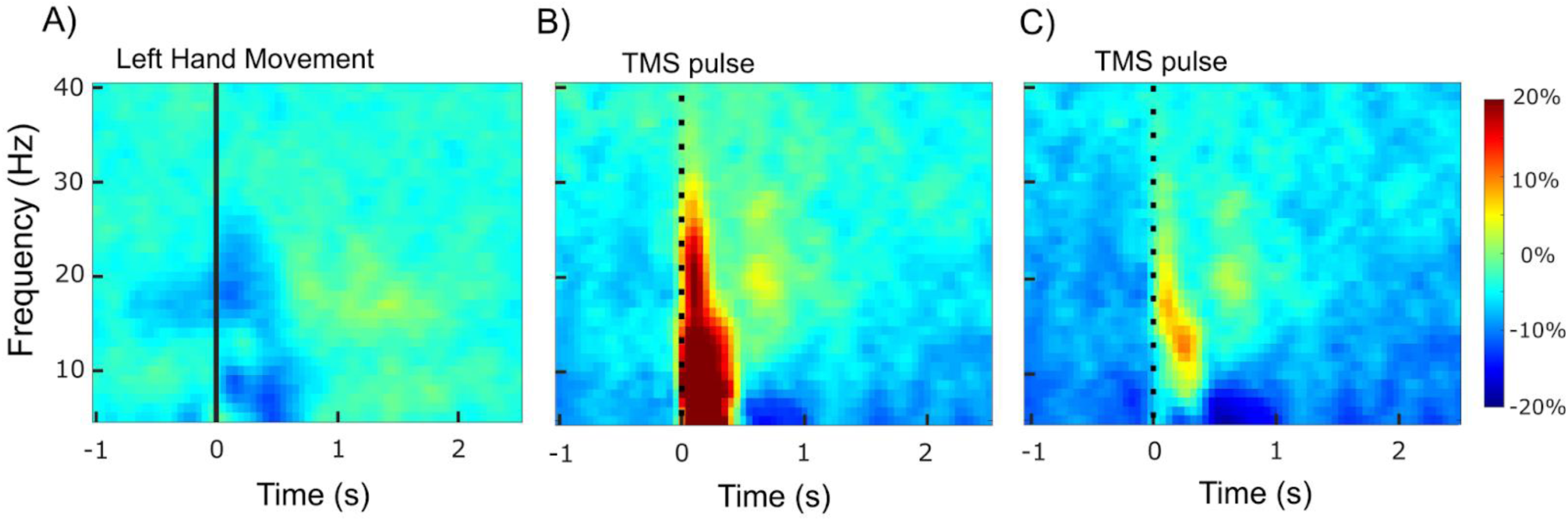
Group-averaged time-frequency map of the right First Dorsal Interosseous Electromyography amplitude during the left hand movement task (A), the ipsilateral Silent Period task prior to subtracting the averaged iSP (B), and the iSP task after subtracting the averaged iSP (C). The vertical black lines indicate the hand movement onset or Transcranial Magnetic Stimulation (TMS) pulse onset, respectively.

Figure 4B presents the group-averaged time-frequency map of the amplitude of the rFDI EMG signal in the iSP task. Contrasting with Figure 4A, it shows a large-scale enhancement of spectral amplitude at frequencies up to 30 Hz after TMS pulse, along with a discernible maximum at beta frequencies localized between 50 and 200 ms. In the time interval from –500 to 1000 ms, EMG *β_size_* was 42.1 ± 31.0 time-frequency bins and was statistically significant in 32 (80 %) participants (all *p* < 0.05).

Figure 4C presents the same map obtained after subtracting each subject’s averaged unrectified EMG response, thereby focusing only on the induced response. Removal of the averaged EMG resulted in a broad-band attenuation of amplitude in the 5–30 Hz range, showing that a TMS-evoked response contributed largely to the initial amplitude modulation. The TMS-induced enhancement of amplitude was restricted to the low-beta frequency range. In the time interval from –500 to 1000 ms, this induced EMG βsize was 27.48 ± 25.5 time-frequency bins, which was statistically significant in 25 (63 %) participants (all *p* < 0.05).

### 3.4 Beta burst characteristics

Figure 4 presents the evolution of beta bursts’ characteristics (probability of occurrence and duration) as derived from the right rFDI EMG relative to movement onset or TMS stimulation. Compared to baseline, the probability of observing a beta burst was significantly reduced in the time windows centered from 0 ms to 100 ms relative to hand movement (peak at the 0 ms time window, *t*_39_ = 3.13, *p_uncorrected_* = 0.0033, *p_corrected_* = 0.0197) and significantly increased in the time windows centered from 0 to 200 ms relative to an ipsilateral TMS pulse (peak at the 0 ms time window, *t*_39_ = -9.69, *p_uncorrected_* < 0.00001, *p_corrected_* < 0.00001; Figure 5A). The average beta burst duration was not significantly modulated during the hand movement task (*p_uncorrected_* > 0.05), and was significantly increased in the time windows centered from 100 to 200 ms relative to the ipsilateral TMS pulse (peak at the 100 ms time window, *t*_39_ = -4.67, *p_uncorrected_* = 0.00004, *p_corrected_* = 0.0002; Figure 5B). Therefore, the large-scale amplitude suppression during movement execution (Figure 4A) reflects the suppression of beta bursts and the large-scale amplitude enhancement following an iSP-inducing TMS stimulation (Figure 4B and 4C) reflects TMS-evoked beta bursts.

**Figure 5.**
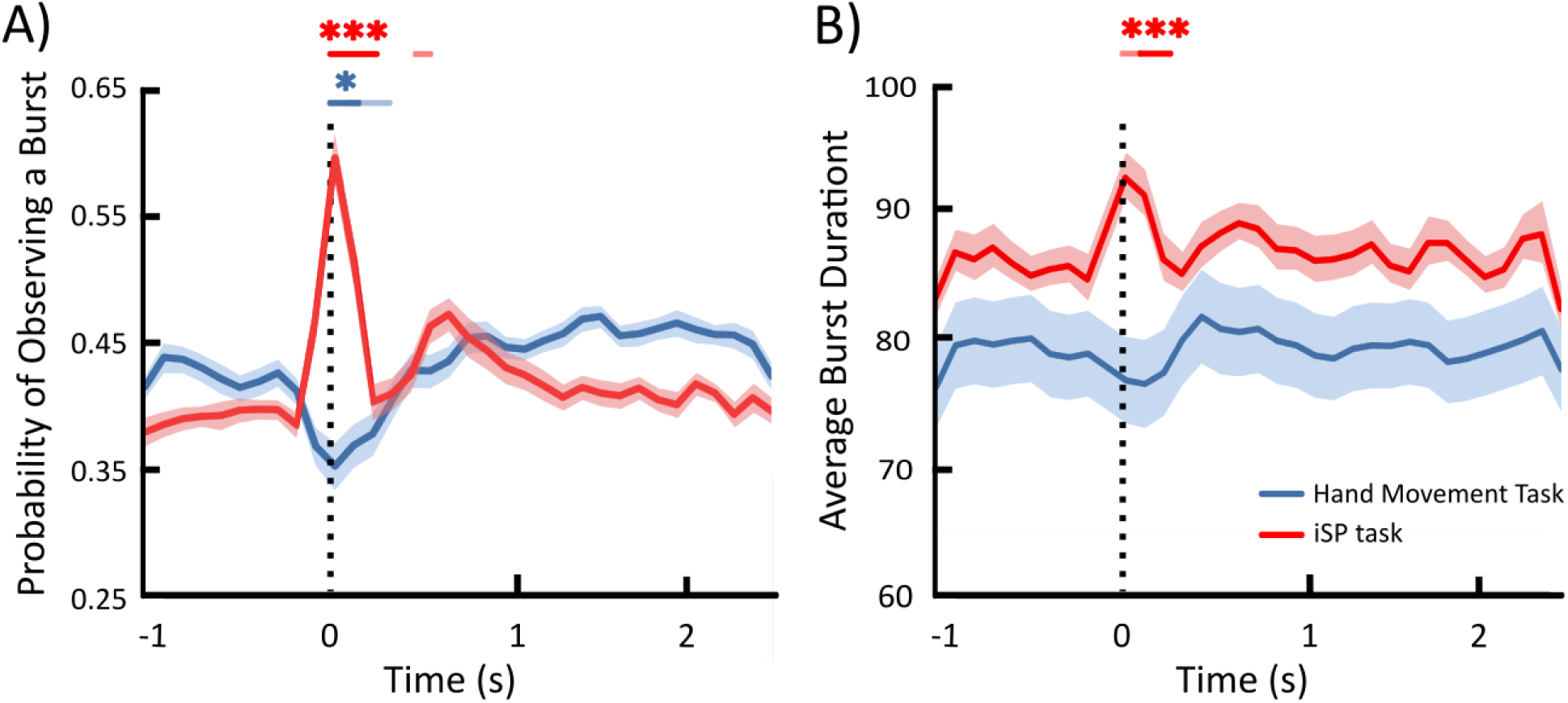
Probability of occurrence in a 200-ms time window (A) and average duration (B) of beta bursts detected in the Electromyography signal from the hand movement execution and ipsilateral Silent Period (iSP) task. Full traces represent the mean across participants and shaded areas indicate standard errors of the mean. The vertical dotted black line indicates the hand movement onset or Transcranial Magnetic Stimulation pulse onset, respectively. Colored lines above the main figure represent time points in which Probability of burst occurrence or average burst duration deviated significantly from baseline before (non-bold) and after (bold) Bonferroni correction. Asterisks represent significance levels with: *, p < 0.05; and ***, p < 0.001

### 3.5 Phase domain EMG changes

For investigating the identified TMS-evoked beta bursts further, Figure 6 presents the time-course of the averaged normalized non-rectified EMG traces filtered through 13– 30 Hz in the TMS-iSP protocol. It highlights the presence of a beta oscillation starting from 20–30 ms after the TMS pulse. Importantly, the first peak of this oscillation appeared consistently at 40 ms across participants. And indeed, the phase consistency analysis revealed a significant phase alignment of Hilbert coefficients in the time interval 10 to 70 ms following the onset of the TMS pulse (all *p* < 0.05), with the beta phase alignment reaching maximal significance at 40 ms after the TMS pulse (*p* = 0.0003; Figure 6). This demonstrates that 10 to 70 ms after the TMS pulse, ipsilateral EMG beta oscillations align in phase and then diverge over time, possibly due to interindividual variability in the shape of the beta bursts.

**Figure 6.**
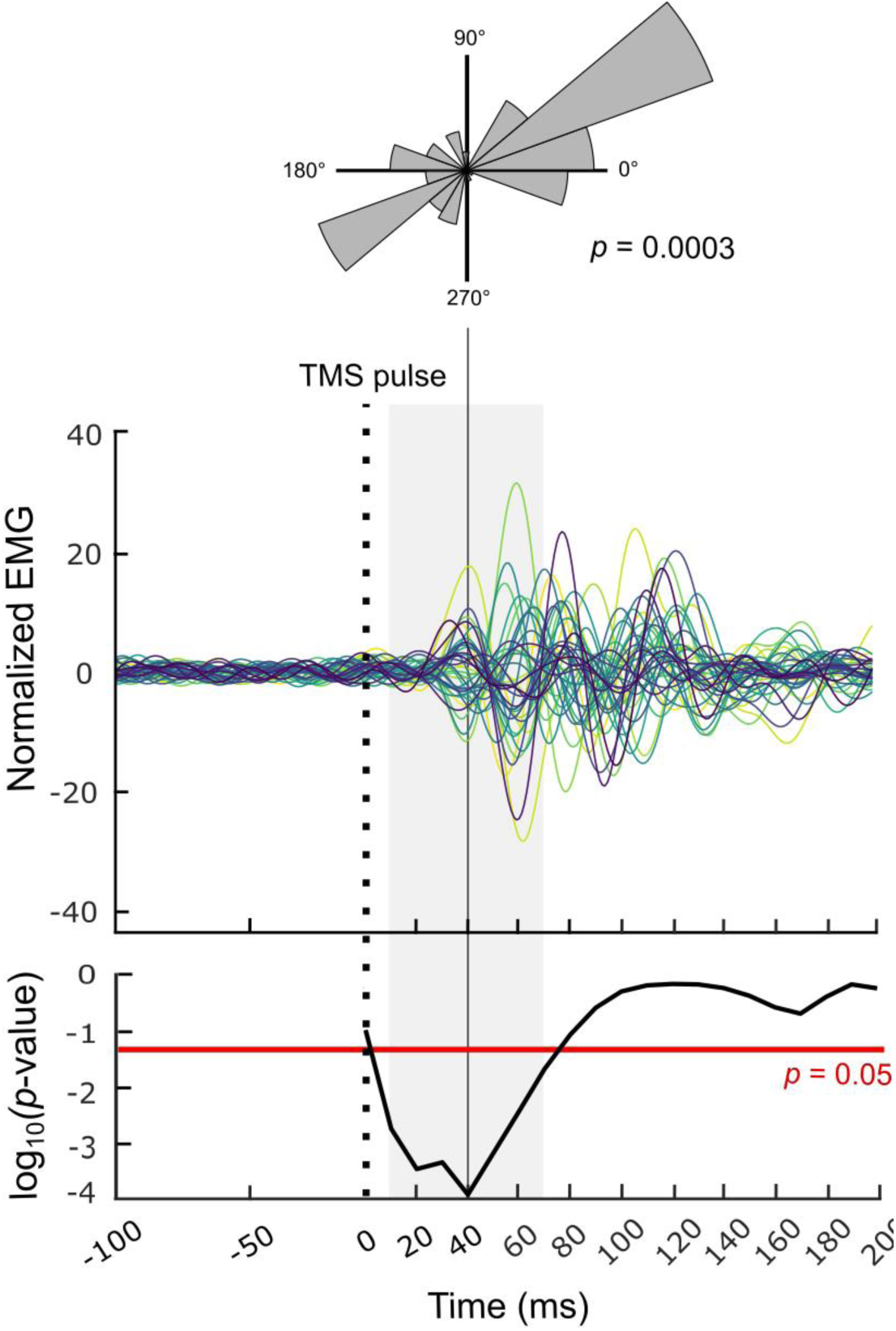
Normalized non-rectified Electromyography (EMG) traces (averaged across trials) of each participant in the ipsilateral Silent Period (iSP) task demonstrating the signals’ phase alignment, along with its corresponding p-value. The vertical dotted black line indicates the Transcranial Magnetic Stimulation (TMS) pulse; the horizontal red line indicates the threshold for statistical significance and the shaded gray area indicates the time period of significant phase alignment. The top inset presents the phase distribution of the Hilbert coefficients across participants. For each angle interval (20 in total, covering 0 to 360°), it presents the sum of Hilbert coefficients’ amplitude whose phase lies in this interval.

To ensure that the observed TMS-evoked beta burst and its phase alignment across participants was not simply a consequence of narrow band-pass filtering, we repeated our analysis on the EMG signals filtered between 31 – 48 Hz. This equally wide frequency band on the neighboring gamma range did not show a significant phase alignment of Hilbert coefficients in the time interval 10 to 70 ms following the onset of the TMS pulse (0.069 ≤ p ≤ 0.99). This suggests that the observed results in the beta band are not simply a consequence of the applied methodology.

### 3.5 Manual Dexterity

Participants scored 15 ± 2 (range 11–19) on *PPT_LH_*, 16 ± 2 (range 10–22) on *PPT_RH_*, 12 ± 2 (range 7–16) on *PPT_BH_*, and 37 ± 7 (range 22–51) on *PPT_Assembly_*. The scores on all PPT subtests are consistent with the findings of other investigators and with PPT population norms from healthy adults (Yeudall et al., 1986; Gu et al., 2023), therefore indicating that our participants completed the PPT adequately.

With respect to the additional bimanual coordination scores, participants scored 2.3 ± 0.4 (range 1.4–3.1) on *PPT_C1_* (PPT*_Assembly_*/(√*PPT_LH_* × √*PPT_RH_*) and 3.1 ± 0.6 (range 2.0– 5.9) on *PPT_C2_* (*PPT_Assembly_*/*PPT_BH_*). Given that twice as much time is allowed for the assembly task, the fact that *PPT_C1_* values were above 2 on average indicates that performance with the two hands working in coordination was better compared to independent performance with each hand individually. Besides, given that *PPT_BH_* indicates a number of peg pairs while *PPT_Assembly_* accounts for each element, a *PPT_C2_* below 4 indicates that performance with the two hands working in coordination was worse compared to performance with the two hands working in synchrony.

### 3.6 Association analyses

The RDA aimed at assessing the associations between the indices of left-/right-hemisphere beta imbalance (Δ*β_size_* and Δ*β_depth_*), iSP (latency, duration, and normalized area), and bimanual motor performance (*PPT_BH_, PPT_Assembly_, PPT_C1_,* and *PPT_C2_*) revealed that, collectively, the beta modulation indices did not explain a significant proportion of variance of the iSP indices (4.4 %, *F*_2,37_ = 0.85, *p* = 0.48). It also revealed that, collectively, the beta modulation and iSP indices did not explain a significant proportion of variance of bimanual dexterity (12.7 %, *F*_5,34_ = 0.99, *p* = 0.44).

The results of a follow-up univariate correlation analysis between all variables are presented in Figure 7. As expected, the contrasts of ipsilateral/contralateral beta imbalance were significantly correlated with each other (*r* = 0.45, *p* = 0.004). The iSP indices were also significantly correlated with each other (0.46 ≤ | *r |* ≤ 0.73, all *p* < 0.05) in the respective expected direction. However, no significant correlation between the Δ*β_size_* and Δ*β_depth_* and measures of the iSP was found (0.26 ≤ *p* ≤ 0.57) and, with the exception of a moderate significant correlation between the Δ*β_size_* and the *PPT_C1_* (*r* = 0.37, *p* = 0.018) no index of left-/right beta imbalance or of iSP significantly correlated with any measure of bimanual motor performance.

**Figure 7.**
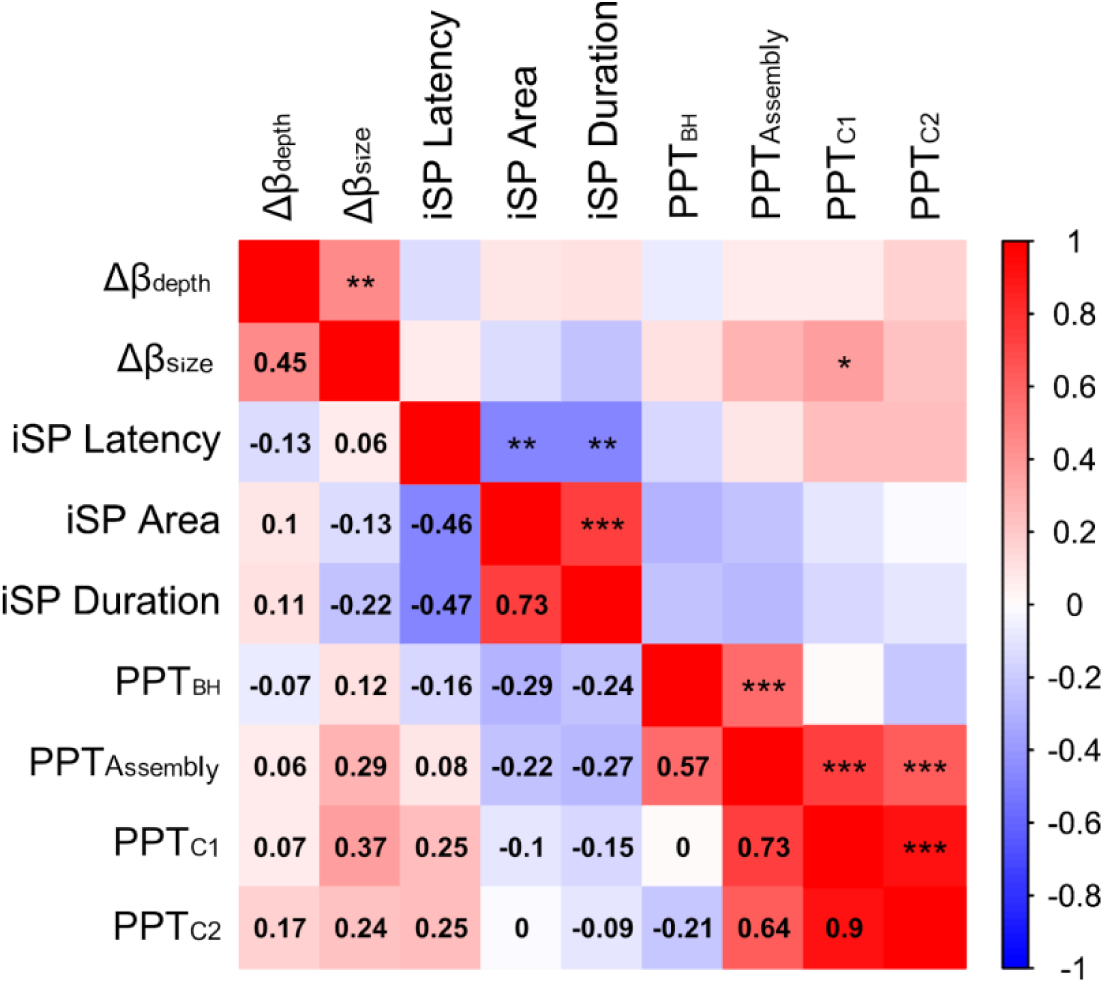
Pairwise correlations between the measures of left-/right-hemisphere beta, the ipsilateral Silent Period (iSP), and Purdue Pegboard Test (PPT). Values below the main diagonal depict rho values, while values above the main diagonal depict the corresponding uncorrected significance levels with: *, p < 0.05; **, p < 0.01; ***, p < 0.001.

## 4. Discussion

Our study aimed at elucidating the link between the movement-related bilateral modulations of SM1 beta oscillations and TMS-induced IHI processes, and investigating the behavioral relevance of the two neurophysiological phenomena. We did not find a significant association across participants in the left-/right-hemisphere imbalance in SM1 beta suppression during volitional hand movement, the iSP, and bimanual dexterity. However, we found that TMS-iSP is contingent on the rapid generation of a phase-aligned beta burst in the EMG proxy signal of the brain hemisphere contralateral to the stimulation, a process exactly opposite in nature to the bilateral beta burst suppression observed during volitional movement.

### 4.1 Contrasting modulation of contralateral SM1 beta oscillations by volitional hand movement and TMS

Using the FDI EMG of the contracted right hand as a proxy for the changes occurring in the contralateral SM1 (Bourguignon et al., 2017; Bräcklein et al., 2022; Echeverria-Altuna et al., 2022; Mongold et al., 2022), we demonstrated a striking divergence in beta SM1 oscillatory dynamics during volitional hand movement and during TMS-evoked iSP. Indeed, ipsilateral SM1 beta bursting activity was suppressed following a hand movement, consistent with an extensive literature (Pfurtscheller and Lopes da Silva, 1999; McFarland et al., 2000; van Wijk et al., 2012; Zaepffel et al., 2013; Kilavik et al., 2013; Démas et al., 2020), while delivering a TMS pulse to SM1 resulted in a significant enhancement of contralateral beta amplitude attributable to the generation of beta bursts. In addition, more pronounced ipsilateral beta amplitude suppression (relative to contralateral) was not significantly associated with classical iSP parameters. Therefore, the ubiquitous beta rhythm suppression—or decrease in beta burst occurrence—in the hemisphere ipsilateral to hand movement appears to receive minimal contribution from the IHI regulatory processes captured by the iSP. Even if naturally occurring IHI processes during volitional movement would encompass a contralateral beta burst generation of the type highlighted in our iSP results, the presence of such a hypothetical burst would be obscured by the co-occurring large-scale beta suppression. In any case, the absence of significant association between measures of movement-related beta modulation and TMS-evoked iSP suggests that these two phenomena reflect two distinct types of interhemispheric interactions occurring at different time scales.

The distinct nature of the beta rhythm changes during movement compared to TMS-iSP can be explained by the vastly distinct state of the motor system during each task (Fuggetta et al., 2005). The suppression in the EEG beta oscillations during volitional movements results from goal-directed action planning, followed by motor command formulation, initiation, and constant refinement, as orchestrated by a distributed prefrontal (Andersen and Cui, 2009), premotor (Halsband et al., 1993), SM1, and parietal (Riemann and Lephart, 2002) system. On the other hand, the IHI processes captured by iSP are manifestations of a very rapid and strong excitation of an SM1 hotspot in the absence of any volition to move. Such excitation does not have the crucial role of precisely executing a desired movement and therefore engages the distributed sensorimotor system in a vastly distinct way. Therefore, in the case of TMS-iSP, the TMS-evoked beta burst in the contralateral hemisphere causes a fleeting inhibitory process which interacts with the natural brain oscillations, yet lacks the functional relevance of the amplitude modulations of the beta rhythm during movement execution. Overall, these considerations suggest that TMS protocols aimed at assessing IHI indeed capture an inhibitory mechanisms across the two SM1s, however, this mechanism is only a restricted part of the complex neurophysiology of the interhemispheric signaling which realizes—and assures the precision of—volitional everyday movement.

### 4.2 TMS-evoked phase resetting of contralateral SM1 beta oscillations

We found that the beta enhancement observed in the proxy signal for the SM1 contralateral to that stimulated with TMS coincides with a consistent phase alignment of beta oscillations both within and across participants. This alignment was most evident ∼40 ms after the TMS pulse in the proxy rFDI EMG signal, but would likely occur ∼20 ms after TMS in contralateral SM1 given the well-documented ∼20-ms propagation delay from SM1 to hand EMG (Ibáñez et al., 2021). The beta phase alignment indicates that a phase-resetting mechanism underlies the iSP. Phase resetting effects of TMS have previously been reported for naturally occurring theta, alpha, and beta oscillatory activity within the stimulated SM1 (Kawasaki et al., 2014; Premoli et al., 2017), but not in the contralateral SM1 and not in relation to IHI. Expanding upon the perspective whereby beta oscillations consist of short-lived bursts (Shin et al., 2017; Bonaiuto et al., 2021; Enz et al., 2021; Szul et al., 2023), which have an inhibitory effect on the motor system, our data suggests that it is the phase of the initial peak of the burst that carries the inhibitory effect. This is evidenced by the near-exact co-occurrence of the initial peak of the beta bursts (Figure 6) and the onset of the iSP (Figure 3) at ∼40 ms after the TMS pulse. Although generated by TMS, the identified beta bursts are comparable in duration (∼160 ms; Figure 6) to naturally occurring beta bursts documented with MEG (∼150 ms reported by Law et al., 2022; ∼167 ms reported by Shin et al., 2017; and ∼180 ms reported by Szul et al., 2023) and with Electrocorticography (between ∼150 and ∼200 ms reported by West et al., 2023), however, precise comparison is made difficult by the heterogeneity in methods used to detect bursts and criteria used to define burst onset and offset. Still an interesting difference in the timing of the behavioral inhibitory effect between our TMS-generated bursts and naturally occurring bursts emerges, with our bursts having an inhibitory effect only between ∼40 and 60 ms after their onset and naturally occurring bursts having a much longer inhibitory effect on behavior lasting for 200–400 ms after their onset (Law et al., 2021; Shin et al., 2017). These differences in behavioral inhibition can potentially be accounted for by different burst properties as well as different task demands. First, there is evidence that naturally occurring beta bursts, although quite diverse in shape, can be classified into motifs and that different shape motifs can potentially result in different inhibitory patterns (Rayson et al., 2023; Szul et al., 2023). The TMS-generated bursts could form a shape motif that is distinct from the motifs of naturally occurring beta bursts and thus likewise result in a different inhibition at the behavioral level. Tentative support for this comes from the noticeable difference in peak shape between our TMS-generated bursts (Figure 6) and the Wavelet-like naturally occurring beta bursts (Bonaiuto et al., 2021; Rayson et al., 2023), however, this is to be verified by future TMS-neuroimaging studies with unified methodology for extracting beta burst properties. Furthermore, although the exact neurophysiology of TMS-IHI remains unknown (Ziemann, 2013), a difference in beta shape motifs could result from the TMS procedure recruiting GABA-ergic inhibitory mechanisms distinct from the GABA-ergic thalamo-cortical and within-cortical signalling believed to generate naturally occurring beta bursts (Bonaiuto et al.,2021; Law et al., 2021) and thus have a different inhibitory outcome. Second, the behavioral inhibitory effect of beta bursts can differ between purely motor tasks—like the constant muscle contraction applied in our experiment—and tasks that involve both cognitive and motor components—like the reaction time task applied in other investigations. Accordingly, in the latter it could be the case that beta bursts have a comparable transient ∼40 to 60 ms inhibitory effect on cognitive processing, but that this keeps impacting motor behavior for a longer time.

Therefore, the TMS-iSP protocol we present, able to generate contralateral beta bursts, could serve future investigations of the nature of the beta oscillations by comparing the properties of the TMS-generated burst to naturally-occurring bursts recorded with EEG/MEG as well as to the predictions of computational models of neural oscillations (Law et al., 2022; West et al., 2023). Additionally, our results provide a promising avenue for developing novel neuromodulation protocols. Those are particularly important considering the often documented abnormalities of beta oscillations in disorders such as cerebral palsy (Démas et al., 2019), multiple sclerosis (Arpin et al., 2017); stroke (Shiner et al., 2015), and Alzheimer’s diseases (de Haan et al., 2008), in all of which a pattern of pathologically suppressed beta burst amplitude is associated with worse clinical outcomes. Therefore, the TMS-iSP protocol could be used in a biofeedback setup for non-invasive modulatory enhancement of beta oscillatory activity, which could lead to therapeutic effects.

### 4.3 Limitations and perspectives

Since in our study we assessed cortical beta oscillations, iSP, and bimanual dexterity non-simultaneously, we could only capitalize on inter-individual variability to identify brain-behavioral relationships The absence of significant association of beta suppression and iSP magnitudes with bimanual dexterity appears puzzling as the PPT is a valid test specifically designed to assess bimanual motor coordination (Lindstrom-Hazel & VanderVlies Veenstra, 2015) and successfully applied in numerous contemporary investigations (e.g., Gu et al., 2023; Hidese et al., 2018; Hinkle & Pontone., 2021; Mongold et al., 2024). Moreover, our participants achieved scores consistent with the normative data and with the results of other investigators (Gu et al., 2023), while displaying a substantial amount of performance variability that could, in theory, have been accounted for by the differences in bilateral beta oscillatory activity or the iSP. However, previous research has found behavioral relevance of bilateral beta modulation and iSP predominantly with in-task assessments of motor performance (Murase et al., 2004; Kuo et al., 2019; Sugata et al., 2020; Morishita et al., 2022) while investigations quantifying motor performance with separately performed tests such as the PPT paint a less clear picture (Gu et al., 2023). Future studies should clarify the task-specificity of the behavioral relevance of beta oscillations and IHI. If beta oscillations and IHI truly are the crucial motor system regulatory mechanisms they are theorized to be (Perez and Cohen, 2009; Engel and Fries, 2010), their role for motor behavior should likely extend beyond in-task effects.

Our investigation focused on a sample of right-handed participants and investigated IHI from the non-dominant towards the dominant hemisphere. However, there is evidence that left-handed participants have more efficient interhemispheric communication (Cherbuin & Brinkman, 2006, but see Raaf & Westerhausen, 2023 for a cautionary note on the matter), and that IHI tends to be stronger from the dominant towards the non-dominant hemisphere (Bäumer et al., 2007; Duque et al., 2007). Therefore, the results might have been more salient (i.e., the transcallosally generated beta bursts would be larger) if the TMS pulse had been applied over the dominant hemisphere. This should be verified in future studies applying our protocol in a mixed sample of left- and right handed participants, while also manipulating the laterality of the TMS pulse.

### 4.4 Conclusion

Our study aimed at clarifying the beta oscillatory dynamics underlying TMS IHI, using a paradigm in which beta oscillations and IHI were assessed non-simultaneously. We found that the large-scale cortical beta amplitude suppression observed during bodily movement was not significantly associated with IHI assessed with the iSP. Moreover, we show that the iSP is contingent on the rapid generation of a high-amplitude phase-aligned beta burst in the hemisphere contralateral to the stimulation, a process opposite in nature to the ipsilateral beta burst suppression observed during movement. Therefore, the phenomena observed in TMS-IHI studies may not generalize to natural motor control, however, the TMS iSP protocol offers a means to evoke and analyze beta bursts, as well as to support neuroimaging and computational modeling studies.

## Ethics Statement

The experimental protocol was approved by the ethics committee of the CUB Hôpital Erasme and all participants signed a detailed informed consent prior to participation.

## Data and Code Availability

All data and code are publicly available on the Open Science Framework (https://osf.io/d5m74/).

## CRediT Author contribution

**Christian Georgiev**: Conceptualization, Investigation, Formal analysis, Visualization, Writing – original draft. **Scott J. Mongold**: Conceptualization, Investigation, Writing – review & editing. **Pierre Cabaraux**: Investigation, Writing – review & editing. **Gilles Naeije**: Writing – review & editing. **Julie Duqué:** Writing – review & editing. **Mathieu Bourguignon**: Conceptualization, Formal analysis, Visualization, Data curation, Supervision, Writing - review & editing.

## References

Andersen, R. A., & Cui, H. (2009). Intention, action planning, and decision making in parietal-frontal circuits. Neuron, 63(5), 568–583. 10.1016/j.neuron.2009.08.028

Arpin, D. J., Heinrichs-Graham, E., Gehringer, J. E., Zabad, R., Wilson, T. W., & Kurz, M. J. (2017). Altered sensorimotor cortical oscillations in individuals with multiple sclerosis suggests a faulty internal model. Human Brain Mapping, 38(8), 4009– 4018. 10.1002/hbm.23644

Bäumer, T., Dammann, E., Bock, F., Klöppel, S., Siebner, H. R., & Münchau, A. (2007). Laterality of interhemispheric inhibition depends on handedness. Experimental Brain Research, 180(2), 195–203. 10.1007/s00221-007-0866-7

Bigdely-Shamlo, N., Mullen, T., Kothe, C., Su, K. M., & Robbins, K. A. (2015). The PREP pipeline: standardized preprocessing for large-scale EEG analysis. Frontiers in Neuroinformatics, 9, 16. 10.3389/fninf.2015.00016

Bologna, M., Caronni, A., Berardelli, A., & Rothwell, J. C. (2012). Practice-related reduction of electromyographic mirroring activity depends on basal levels of interhemispheric inhibition. The European Journal of Neuroscience, 36(12), 3749– 3757. 10.1111/ejn.12009

Bonaiuto, J. J., Little, S., Neymotin, S. A., Jones, S. R., Barnes, G. R., & Bestmann, S. (2021). Laminar dynamics of high amplitude beta bursts in human motor cortex. NeuroImage, 242, 118479. 10.1016/j.neuroimage.2021.118479

Bourguignon, M., Piitulainen, H., Smeds, E., Zhou, G., Jousmäki, V., & Hari, R. (2017). MEG Insight into the spectral dynamics underlying steady isometric muscle contraction. The Journal of neuroscience : the official journal of the Society for Neuroscience, 37(43), 10421–10437. 10.1523/JNEUROSCI.0447-17.2017

Bräcklein, M., Barsakcioglu, D. Y., Del Vecchio, A., Ibáñez, J., & Farina, D. (2022). Reading and Modulating Cortical β Bursts from Motor Unit Spiking Activity. The Journal of neuroscience : the official journal of the Society for Neuroscience, 42(17), 3611–3621. 10.1523/JNEUROSCI.1885-21.2022

Buzsáki, G., & Draguhn, A. (2004). Neuronal oscillations in cortical networks. Science, 304(5679), 1926–1929. 10.1126/science.1099745

Canavier C. C. (2015). Phase-resetting as a tool of information transmission. Current Opinion in Neurobiology, 31, 206–213. 10.1016/j.conb.2014.12.003

Cherbuin, N., & Brinkman, C. (2006). Hemispheric interactions are different in left-handed individuals. Neuropsychology, 20(6), 700–707. 10.1037/0894-4105.20.6.700

de Haan, W., Stam, C. J., Jones, B. F., Zuiderwijk, I. M., van Dijk, B. W., & Scheltens, P. (2008). Resting-state oscillatory brain dynamics in Alzheimer disease. Journal of clinical neurophysiology, 25(4), 187–193. 10.1097/WNP.0b013e31817da184

Démas, J., Bourguignon, M., Périvier, M., De Tiège, X., Dinomais, M., & Van Bogaert, P. (2020). Mu rhythm: State of the art with special focus on cerebral palsy. Annals of Physical and Rehabilitation Medicine, 63(5), 439–446. 10.1016/j.rehab.2019.06.007

Duque, J., Mazzocchio, R., Dambrosia, J., Murase, N., Olivier, E., & Cohen, L. G. (2005). Kinematically specific interhemispheric inhibition operating in the process of generation of a voluntary movement. Cerebral Cortex, 15(5), 588–593. 10.1093/cercor/bhh160

Duque, J., Murase, N., Celnik, P., Hummel, F., Harris-Love, M., Mazzocchio, R., Olivier, E., & Cohen, L. G. (2007). Intermanual Differences in movement-related interhemispheric inhibition. Journal of Cognitive Neuroscience, 19(2), 204–213. 10.1162/jocn.2007.19.2.204

Echeverria-Altuna, I., Quinn, A. J., Zokaei, N., Woolrich, M. W., Nobre, A. C., & van Ede, F. (2022). Transient beta activity and cortico-muscular connectivity during sustained motor behaviour. Progress in Neurobiology, 214, 102281. 10.1016/j.pneurobio.2022.102281

Engel, A. K., & Fries, P. (2010). Beta-band oscillations--signalling the status quo?. Current Opinion in Neurobiology, 20(2), 156–165. 10.1016/j.conb.2010.02.015

Enz, N., Ruddy, K. L., Rueda-Delgado, L. M., & Whelan, R. (2021). Volume of β-Bursts, But Not Their Rate, Predicts Successful Response Inhibition. The Journal of Neuroscience: the official journal of the Society for Neuroscience, 41(23), 5069– 5079. 10.1523/JNEUROSCI.2231-20.2021

Erickson, B., Kim, B., Sabes, P., Rich, R., Hatcher, A., Fernandez-Nuñez, G., Mentzelopoulos, G., Vitale, F., & Medaglia, J. (2024). TMS-induced phase resets depend on TMS intensity and EEG phase. Journal of Neural Engineering, 21(5), 056035. 10.1088/1741-2552/ad7f87

Ferbert, A., Priori, A., Rothwell, J. C., Day, B. L., Colebatch, J. G., & Marsden, C. D. (1992). Interhemispheric inhibition of the human motor cortex. The Journal of physiology, 453, 525–546. 10.1113/jphysiol.1992.sp019243

Fleming, M. K., & Newham, D. J. (2017). Reliability of Transcallosal Inhibition in Healthy Adults. Frontiers in human neuroscience, 10, 681. 10.3389/fnhum.2016.00681

Fling, B. W., Benson, B. L., & Seidler, R. D. (2013). Transcallosal sensorimotor fiber tract structure-function relationships. Human Brain Mapping, 34(2), 384–395. 10.1002/hbm.21437

Fuggetta, G., Fiaschi, A., & Manganotti, P. (2005). Modulation of cortical oscillatory activities induced by varying single-pulse transcranial magnetic stimulation intensity over the left primary motor area: a combined EEG and TMS study. NeuroImage, 27(4), 896–908. 10.1016/j.neuroimage.2005.05.013

Garvey, M. A., Ziemann, U., Becker, D. A., Barker, C. A., & Bartko, J. J. (2001). New graphical method to measure silent periods evoked by transcranial magnetic stimulation. Clinical Neurophysiology: official journal of the International Federation of Clinical Neurophysiology, 112(8), 1451–1460. 10.1016/s1388-2457(01)00581-8

Giovannelli, F., Borgheresi, A., Balestrieri, F., Zaccara, G., Viggiano, M. P., Cincotta, M., & Ziemann, U. (2009). Modulation of interhemispheric inhibition by volitional motor activity: an ipsilateral silent period study. The Journal of Physiology, 587(Pt 22), 5393–5410. 10.1113/jphysiol.2009.175885

Gu, B., Wang, K., Chen, L., He, J., Zhang, D., Xu, M., Wang, Z., & Ming, D. (2023). Study of the Correlation between the Motor Ability of the Individual Upper Limbs and Motor Imagery Induced Neural Activities. Neuroscience, 530, 56–65. 10.1016/j.neuroscience.2023.08.032

Halsband, U., Ito, N., Tanji, J., & Freund, H. J. (1993). The role of premotor cortex and the supplementary motor area in the temporal control of movement in man. Brain: a journal of neurology, 116 ( *Pt 1*), 243–266. 10.1093/brain/116.1.243

Heinrichs-Graham, E., Kurz, M. J., Gehringer, J. E., & Wilson, T. W. (2017). The functional role of post-movement beta oscillations in motor termination. Brain structure & function, 222(7), 3075–3086. 10.1007/s00429-017-1387-1

Hidese, S., Ota, M., Sasayama, D., Matsuo, J., Ishida, I., Hiraishi, M., Teraishi, T., Hattori, K., & Kunugi, H. (2018). Manual dexterity and brain structure in patients with schizophrenia: A whole-brain magnetic resonance imaging study. Psychiatry research. Neuroimaging, 276, 9–14. 10.1016/j.pscychresns.2018.04.003

Hinkle, J. T., & Pontone, G. M. (2021). Psychomotor processing and functional decline in Parkinson’s disease predicted by the Purdue Pegboard test. International journal of geriatric psychiatry, 36(6), 909–916. 10.1002/gps.5492

Hupfeld, K. E., Swanson, C. W., Fling, B. W., & Seidler, R. D. (2020). TMS-induced silent periods: A review of methods and call for consistency. Journal of neuroscience methods, 346, 108950. 10.1016/j.jneumeth.2020.108950

Hussain, S. J., Cohen, L. G., & Bönstrup, M. (2019). Beta rhythm events predict corticospinal motor output. Scientific reports, 9(1), 18305. 10.1038/s41598-019-54706-w

Hyvärinen, A., & Oja, E. (2000). Independent component analysis: algorithms and applications. Neural networks: the official journal of the International Neural Network Society, 13(4-5), 411–430. 10.1016/s0893-6080(00)00026-5

Ibáñez, J., Del Vecchio, A., Rothwell, J. C., Baker, S. N., & Farina, D. (2021). Only the Fastest Corticospinal Fibers Contribute to β Corticomuscular Coherence. The Journal of neuroscience : the official journal of the Society for Neuroscience, 41(22), 4867–4879. 10.1523/JNEUROSCI.2908-20.2021

Illman, M., Laaksonen, K., Jousmäki, V., Forss, N., & Piitulainen, H. (2022). Reproducibility of Rolandic beta rhythm modulation in MEG and EEG. Journal of neurophysiology, 127(2), 559–570. 10.1152/jn.00267.2021

Jung, P., & Ziemann, U. (2006). Differences of the ipsilateral silent period in small hand muscles. Muscle & Nerve, 34(4), 431–436. 10.1002/mus.20604

Kawasaki, M., Uno, Y., Mori, J., Kobata, K., & Kitajo, K. (2014). Transcranial magnetic stimulation-induced global propagation of transient phase resetting associated with directional information flow. Frontiers in Human Neuroscience, 8, 173. 10.3389/fnhum.2014.00173

Khanna, P., & Carmena, J. M. (2017). Beta band oscillations in motor cortex reflect neural population signals that delay movement onset. eLife, 6, e24573. 10.7554/eLife.24573

Kilavik, B. E., Zaepffel, M., Brovelli, A., MacKay, W. A., & Riehle, A. (2013). The ups and downs of β oscillations in sensorimotor cortex. Experimental Neurology, 245, 15–26. 10.1016/j.expneurol.2012.09.014

Kuo, Y. L., Dubuc, T., Boufadel, D. F., & Fisher, B. E. (2017). Measuring ipsilateral silent period: Effects of muscle contraction levels and quantification methods. Brain Research, 1674, 77–83. 10.1016/j.brainres.2017.08.015

Kuo, Y. L., Kutch, J. J., & Fisher, B. E. (2019). Relationship between Interhemispheric Inhibition and Dexterous Hand Performance in Musicians and Non-musicians. Scientific Reports, 9(1), 11574. 10.1038/s41598-019-47959-y

Law, R. G., Pugliese, S., Shin, H., Sliva, D. D., Lee, S., Neymotin, S., Moore, C., & Jones, S. R. (2022). Thalamocortical mechanisms regulating the relationship between transient beta events and human tactile perception. Cerebral cortex, 32(4), 668–688. 10.1093/cercor/bhab221

Lindstrom-Hazel, D. K., & VanderVlies Veenstra, N. (2015). Examining the Purdue Pegboard Test for occupational therapy practice. The Open Journal of Occupational Therapy, 3(3). 10.15453/2168-6408.1178

Mary, A., Wens, V., Op de Beeck, M., Leproult, R., De Tiège, X., & Peigneux, P. (2017). Age-related differences in practice-dependent resting-state functional connectivity related to motor sequence learning. Human brain mapping, 38(2), 923–937. 10.1002/hbm.23428

McFarland, D. J., Miner, L. A., Vaughan, T. M., & Wolpaw, J. R. (2000). Mu and beta rhythm topographies during motor imagery and actual movements. Brain topography, 12(3), 177–186. 10.1023/a:1023437823106

Mongold, S. J., Piitulainen, H., Legrand, T., Vander Ghinst, M., Naeije, G., Jousmäki, V., & Bourguignon, M. (2022). Temporally stable beta sensorimotor oscillations and corticomuscular coupling underlie force steadiness. NeuroImage, 261, 119491. 10.1016/j.neuroimage.2022.119491.

Mongold, S. J., Georgiev, C., Legrand, T., & Bourguignon, M. (2024). Afferents to action: Cortical proprioceptive processing assessed with corticokinematic coherence Specifically Relates to Gross Motor Skills. eNeuro, 11(1), ENEURO.0384-23.2023. 10.1523/ENEURO.0384-23.2023

Morishita, T., Timmermann, J. E., Schulz, R., & Hummel, F. C. (2022). Impact of interhemispheric inhibition on bimanual movement control in young and old. Experimental Brain Research, 240(2), 687–701. 10.1007/s00221-021-06258-7

Murase, N., Duque, J., Mazzocchio, R., & Cohen, L. G. (2004). Influence of interhemispheric interactions on motor function in chronic stroke. Annals of Neurology, 55(3), 400–409. 10.1002/ana.10848

Neuper, C., Wörtz, M., & Pfurtscheller, G. (2006). ERD/ERS patterns reflecting sensorimotor activation and deactivation. Progress in brain research, 159, 211–222. 10.1016/S0079-6123(06)59014-4

O’Keeffe, A. B., Malekmohammadi, M., Sparks, H., & Pouratian, N. (2020). Synchrony drives motor cortex beta bursting, waveform dynamics, and phase-amplitude coupling in Parkinson’s disease. The Journal of neuroscience, 40(30), 5833–5846. 10.1523/JNEUROSCI.1996-19.2020

Oldfield R. C. (1971). The assessment and analysis of handedness: the Edinburgh inventory. Neuropsychologia, 9(1), 97–113. 10.1016/0028-3932(71)90067-4

Oostenveld, R., Fries, P., Maris, E., & Schoffelen, J. M. (2011). FieldTrip: Open source software for advanced analysis of MEG, EEG, and invasive electrophysiological data. Computational intelligence and neuroscience, 2011, 156869. 10.1155/2011/156869

Perez, M. A., & Cohen, L. G. (2009). Interhemispheric inhibition between primary motor cortices: what have we learned?. The Journal of physiology, 587(Pt 4), 725–726. 10.1113/jphysiol.2008.166926

Pfurtscheller G. (1981). Central beta rhythm during sensorimotor activities in man. Electroencephalography and clinical neurophysiology, 51(3), 253–264. 10.1016/0013-4694(81)90139-5

Pfurtscheller, G., & Lopes da Silva, F. H. (1999). Event-related EEG/MEG synchronization and desynchronization: basic principles. Clinical neurophysiology : official journal of the International Federation of Clinical Neurophysiology, 110(11), 1842–1857. 10.1016/s1388-2457(99)00141-8

Premoli, I., Bergmann, T. O., Fecchio, M., Rosanova, M., Biondi, A., Belardinelli, P., & Ziemann, U. (2017). The impact of GABAergic drugs on TMS-induced brain oscillations in human motor cortex. NeuroImage, 163, 1–12. 10.1016/j.neuroimage.2017.09.023

Raaf, N., & Westerhausen, R. (2023). Hand preference and the corpus callosum: Is there really no association? Neuroimage Reports, 3(1), 100160. 10.1016/j.ynirp.2023.100160

Rayson, H., Szul, M. J., El-Khoueiry, P., Debnath, R., Gautier-Martins, M., Ferrari, P. F., Fox, N., & Bonaiuto, J. J. (2023). Bursting with Potential: How Sensorimotor Beta Bursts Develop from Infancy to Adulthood. The Journal of Neuroscience, 43(49), 8487–8503. 10.1523/JNEUROSCI.0886-23.2023

Riemann, B. L., & Lephart, S. M. (2002). The Sensorimotor System, Part II: The Role of Proprioception in Motor Control and Functional Joint Stability. Journal of athletic training, 37(1), 80–84.

Rossini, P. M., Barker, A. T., Berardelli, A., Caramia, M. D., Caruso, G., Cracco, R. Q., Dimitrijević, M. R., Hallett, M., Katayama, Y., & Lücking, C. H. (1994). Non-invasive electrical and magnetic stimulation of the brain, spinal cord and roots: basic principles and procedures for routine clinical application. Report of an IFCN committee. Electroencephalography and clinical neurophysiology, 91(2), 79–92. 10.1016/0013-4694(94)90029-9

Salenius, S., & Hari, R. (2003). Synchronous cortical oscillatory activity during motor action. Current opinion in neurobiology, 13(6), 678–684. 10.1016/j.conb.2003.10.008

Sherman, M. A., Lee, S., Law, R., Haegens, S., Thorn, C. A., Hämäläinen, M. S., Moore, C. I., & Jones, S. R. (2016). Neural mechanisms of transient neocortical beta rhythms: Converging evidence from humans, computational modeling, monkeys, and mice. Proceedings of the National Academy of Sciences of the United States of America, 113(33), E4885–E4894. 10.1073/pnas.1604135113

Shin, H., Law, R., Tsutsui, S., Moore, C. I., & Jones, S. R. (2017). The rate of transient beta frequency events predicts behavior across tasks and species. eLife, 6, e29086. 10.7554/eLife.29086

Shiner, C. T., Tang, H., Johnson, B. W., & McNulty, P. A. (2015). Cortical beta oscillations and motor thresholds differ across the spectrum of post-stroke motor impairment, a preliminary MEG and TMS study. Brain Research, 1629, 26–37. 10.1016/j.brainres.2015.09.037

Sugata, H., Yagi, K., Yazawa, S., Nagase, Y., Tsuruta, K., Ikeda, T., Nojima, I., Hara, M., Matsushita, K., Kawakami, K., & Kawakami, K. (2020). Role of beta-band resting-state functional connectivity as a predictor of motor learning ability. NeuroImage, 210, 116562. 10.1016/j.neuroimage.2020.116562

Szul, M. J., Papadopoulos, S., Alavizadeh, S., Daligaut, S., Schwartz, D., Mattout, J., & Bonaiuto, J. J. (2023). Diverse beta burst waveform motifs characterize movement-related cortical dynamics. Progress in neurobiology, 228, 102490. 10.1016/j.pneurobio.2023.102490

Tiffin, J., & Asher, E. J. (1948). The Purdue pegboard; norms and studies of reliability and validity. The Journal of applied psychology, 32(3), 234–247. 10.1037/h0061266

Tinkhauser, G., Pogosyan, A., Little, S., Beudel, M., Herz, D. M., Tan, H., & Brown, P. (2017). The modulatory effect of adaptive deep brain stimulation on beta bursts in Parkinson’s disease. Brain: a journal of neurology, 140(4), 1053–1067. 10.1093/brain/awx010

Van Den Wollenberg, A. L. (1977). Redundancy analysis an alternative for canonical correlation analysis. Psychometrika, 42(2), 207–219. 10.1007/bf02294050

van Wijk, B. C., Beek, P. J., & Daffertshofer, A. (2012). Differential modulations of ipsilateral and contralateral beta (de)synchronization during unimanual force production. The European Journal of Neuroscience, 36(1), 2088–2097. 10.1111/j.1460-9568.2012.08122.x

Vigário, R., Särelä, J., Jousmäki, V., Hämäläinen, M., & Oja, E. (2000). Independent component approach to the analysis of EEG and MEG recordings. IEEE transactions on bio-medical engineering, 47(5), 589–593. 10.1109/10.841330

Voloh, B., & Womelsdorf, T. (2016). A Role of phase-resetting in coordinating large scale neural networks during attention and goal-directed behavior. Frontiers in Systems Neuroscience, 10, 18. 10.3389/fnsys.2016.00018

Wassermann, E. M., Fuhr, P., Cohen, L. G., & Hallett, M. (1991). Effects of transcranial magnetic stimulation on ipsilateral muscles. Neurology, 41(11), 1795–1799. 10.1212/wnl.41.11.1795

Wessel J. R. (2020). β-Bursts Reveal the Trial-to-Trial Dynamics of Movement Initiation and Cancellation. The Journal of Neuroscience : the official journal of the Society for Neuroscience, 40(2), 411–423. 10.1523/JNEUROSCI.1887-19.2019

West, T. O., Duchet, B., Farmer, S. F., Friston, K. J., & Cagnan, H. (2023). When do bursts matter in the primary motor cortex? Investigating changes in the intermittencies of beta rhythms associated with movement states. Progress in Neurobiology, 221, 102397. 10.1016/j.pneurobio.2022.102397.

Yeudall, L. T., Fromm, D., Reddon, J. R., & Stefanyk, W. O. (1986). Normative data stratified by age and sex for 12 neuropsychological tests. Journal of Clinical Psychology, 42(6), 918–946. 10.1002/1097-4679(198611)42:6

Zaepffel, M., Trachel, R., Kilavik, B. E., & Brochier, T. (2013). Modulations of EEG beta power during planning and execution of grasping movements. PloS one, 8(3), e60060. 10.1371/journal.pone.0060060

Ziemann U. (2013). Pharmaco-transcranial magnetic stimulation studies of motor excitability. Handbook of Clinical Neurology, 116, 387–397. 10.1016/B978-0-444-53497-2.00032-2

